# Chronic hM4Di-DREADD mediated chemogenetic inhibition of forebrain excitatory neurons in postnatal or juvenile life does not alter adult mood-related behavior

**DOI:** 10.1101/2021.09.19.460973

**Authors:** Praachi Tiwari, Darshana Kapri, Amartya Pradhan, Angarika Balakrishnan, Pratik R. Chaudhari, Vidita A. Vaidya

## Abstract

G-protein coupled receptors (GPCRs) coupled to Gi-signaling, in particular downstream of monoaminergic neurotransmission, are posited to play a key role during developmental epochs (postnatal and juvenile), in shaping the emergence of adult anxio-depressive behaviors and sensorimotor gating. To address the role of Gi-signaling in these developmental windows, we used a CamKIIα-tTA::TRE hM4Di bigenic mouse line to express the hM4Di-DREADD in forebrain excitatory neurons and enhanced Gi-signaling via chronic administration of the DREADD agonist, CNO in the postnatal (PNCNO: postnatal day 2-14) or juvenile (JCNO: postnatal day 28-40) window. We confirmed that the expression of the HA-tagged hM4Di-DREADD was restricted to CamKII-positive neurons in the forebrain, and administration of CNO in postnatal or juvenile windows evoked inhibition in forebrain circuits of the hippocampus and cortex, as indicated by a decline in expression of the neuronal activity marker, c-fos. hM4Di-DREADD mediated inhibition of CamKIIα-positive forebrain excitatory neurons in postnatal or juvenile life did not impact the weight profile of mouse pups, and also did not influence the normal ontogeny of sensory reflexes. Further, postnatal or juvenile hM4Di-DREADD mediated inhibition of CamKIIα-positive forebrain excitatory neurons did not alter anxiety or despair-like behaviors in adulthood, and did not impact sensorimotor gating. Collectively, these results indicate that chemogenetic induction of Gi-signaling in CamKIIα-positive forebrain excitatory neurons in postnatal and juvenile temporal windows does not appear to impinge on the programming of anxio-depressive behaviors in adulthood.

## Introduction

Experiences during early developmental windows play a crucial role in the fine-tuning and shaping of an individual’s behavioral and functional responses in adulthood (Ansorge et al., 2007; Bale et al., 2010; Di Segni et al., 2018; Gross and Hen, 2004; Hensch, 2005). While exposure to early stress and trauma is associated with persistent increases in anxiety and despair-like behavior in preclinical studies (Chen and Baram, 2016; De Melo et al., 2018; Targum and Nemeroff, 2019; Wang et al., 2020), enriched environment exposure (Cymerblit-Sabba et al., 2013; Francis et al., 2002; Kempermann et al., 1997; Ravenelle et al., 2014; Sparling et al., 2018) and high maternal care during these early temporal windows is associated with enhanced stress-coping and resilient behavioral responses (Bagot et al., 2009; Bredy et al., 2003; Champagne et al., 2008). The neurotransmitter, serotonin (5-HT), and signaling via the Gi-coupled 5-HT_1A_ and Gq-coupled 5-HT_2A_ receptors, has been implicated in playing an important role in shaping the development of mood-related behavior (Altieri et al., 2015; Gordon and Hen, 2004; Tiwari et al., 2021). Elevation of 5-HT levels during postnatal life, either via pharmacological blockade or genetic loss of function of the 5-HT transporter, is associated with enhanced anxiety and despair-like behavior that persists across the life-span (Ansorge et al., 2004, 2008; Sarkar et al., 2014). Loss of function of the Gq-coupled 5-HT_2A_ receptor, in particular in the forebrain, is associated with reduced anxiety-like behavior (Weisstaub, 2006), whereas loss of function of the Gi-coupled 5-HT_1A_ receptor during postnatal life, in both forebrain and raphe neurocircuits, has been linked to increased anxiety-like behavior (Gross et al., 2002; Mineur et al., 2014; Richardson-Jones et al., 2011, 2010; Vinkers et al., 2010a). Furthermore, pharmacological blockade of the Gi-coupled 5-HT_1A_ receptor during postnatal life is associated with the emergence of increased anxiety in adulthood (Garcia-Garcia et al., 2014; Sarkar et al., 2014; Vinkers et al., 2010b), whereas pharmacological stimulation of the Gq-coupled 5-HT_2A_ receptors (Sarkar et al., 2014) or enhanced Gq-signaling driven via chemogenetic activation of excitatory forebrain neurons during postnatal windows programs increased anxiety and despair-like behavior in adulthood (Pati et al., 2020).

It has been hypothesized that early stress may shift the balance towards enhanced excitatory Gq-coupled signaling accompanied by a decline in inhibitory Gi-coupled signaling in forebrain neurocircuits, which could contribute to the programing of perturbed anxiety and despair-like behaviors (Lambe et al., 2011; Sumner et al., 2008; Tiwari et al., 2021). Chemogenetic studies indicate that enhanced Gq-signaling in forebrain excitatory neurons during postnatal life programs long-lasting increases in anxiety and despair-like behavior along with disrupted sensorimotor gating (Pati et al., 2020). Several preclinical studies suggest that a loss or reduction in signaling via the Gi-coupled 5-HT_1A_ receptor during the postnatal temporal window enhances anxio-depressive behaviors in adulthood (Gross et al., 2002, 2000; Ramboz et al., 1998; Richardson-Jones et al., 2011, 2010; Vinkers et al., 2010a). However, a recent study indicates that enhanced Gi-signaling driven chemogenetically in prefrontal cortical neurons during postnatal life results in enhanced adult anxiety and despair-like behavior, phenocopying the effects of early stress (Teissier et al., 2019). Clinical evidence based on studies of 5-HT_1A_ receptor binding suggest that Gi-coupled receptors may be associated with resilience to anxiety (Albert et al., 2019; Armbruster et al., 2011; Savitz et al., 2009). Collectively, these reports provide impetus for experiments to test whether perturbation of Gi-signaling in forebrain excitatory neurons during early developmental windows can alter the programming of mood-related behaviors.

Here, we directly addressed the influence of increased Gi-mediated signaling in forebrain excitatory neurons in postnatal and juvenile life in the shaping of anxiety and despair-like behavior, as well as sensorimotor gating responses, in adulthood. We used the Gi-coupled inhibitory (hM4Di) Designer Receptors Exclusively Activated by Designer Drugs (DREADD), which were expressed in CamKIIα-positive forebrain excitatory neurons via a bigenic mouse line (CamKIIα-tTA::TRE hM4Di) (Alexander et al., 2009), and used the DREADD ligand clozapine-N-oxide (CNO; 5 mg/kg) (Roth, 2016) to activate Gi-signaling during postnatal (postnatal day 2-14) and juvenile (postnatal day 28-40) windows followed by behavioral analysis in adulthood. We show that hM4Di-DREADD mediated inhibition of CamKIIα-positive forebrain excitatory neurons in either the postnatal or juvenile temporal windows does not influence anxiety- and despair-like behavior, or sensorimotor gating in adulthood.

## Materials and Methods

### Animals

Bigenic CamKIIa-tTA::TRE-hM4Di mice were used for all experiments. The CamKIIa-tTA transgenic mouse line (Mayford et al., 1996) was a gift from Dr. Christopher Pittenger, Department of Psychiatry, Yale School of Medicine. The TRE-hM4Di mouse line was purchased from Jackson Laboratories, USA (Cat. No. 024114; B6.Cg-Tg(tetO-CHRM4*)2Blr/J; Jackson Laboratories, USA). Bigenic animals were generated for the experiments by mating CamKIIa-tTA::TRE-hM4Di males to CamKIIa-tTA::TRE-hM4Di females. The genotypes were confirmed by PCR-based analysis. All dams were individually housed in separate cages and litter size was restricted to 6-8 pups per litter. All animals were bred and maintained in the Tata Institute of Fundamental Research (TIFR), Mumbai (India), animal house facility on a 12 hour light-dark cycle from 7 am to 7 pm, with *ad libitum* access to food and water. Experimental procedures were carried out as per the guidelines of the Committee for the Purpose of Control and Supervision of Experiments on Animals (CPCSEA), Government of India, and were approved by TIFR animal ethics committee. Care was taken across all experiments to minimize any pain or suffering, and to restrict the number of animals used.

### Drug Treatment Paradigms

For postnatal drug treatments, bigenic CamKIIα-tTA::TRE-hM4Di mouse pups were orally administered either Clozapine-N-oxide (CNO) (Cat no 4963, Tocris, UK; 5 mg/kg in 5% sucrose solution containing 1% DMSO) or vehicle (5% sucrose solution containing 1% DMSO) for thirteen days, commencing from postnatal day 2 (P2) to postnatal day 14 (P14). Post weaning (P24-P27), animals were group housed for three months prior to assessment on behavioral assays. For juvenile drug treatments, bigenic CamKIIα-tTA::TRE-hM4Di mouse pups were weaned between P24-P27, group housed and randomly assigned to either the vehicle or CNO treatment groups. Juvenile bigenic CamKIIα-tTA::TRE-hM4Di mice received either CNO (5 mg/kg in 5% sucrose solution containing 1% DMSO) or vehicle (5% sucrose solution containing 1% DMSO) for thirteen days from P28 to P40. All animals were left undisturbed from P41 for two months prior to subjecting them to behavioral testing. To assess whether postnatal (PNCNO) or juvenile (JCNO) CNO treatment influenced the weight of pups, we carried out an extensive weight profiling across the duration of the PNCNO and JCNO treatment paradigms.

### Western blotting

To assess HA-tagged hM4Di-DREADD expression in the hippocampus and cortex of CamKIIα-tTA::TRE-hM4Di bigenic mice at P7 and P35, we carried out western blotting analysis for the HA antigen. To determine the influence of CNO-mediated activation of the hM4Di-DREADD on neuronal activity marker expression (c-fos), we administered a single dose of CNO (5 mg/kg) or vehicle to CamKIIα-tTA::TRE-hM4Di bigenic mouse pups at P7 and to the juvenile cohort at P35 and sacrificed them thirty minutes post-administration. Hippocampi and cortex tissue were dissected and homogenized in Radioimmunoprecipitation assay (RIPA) buffer (50 nM Tris-Cl (pH 8.0), 5 mM EDTA, 1% NP-40, 150 mM NaCl) using a Dounce homogenizer. Protein concentration was estimated with the Quantipro BCA assay kit (Sigma-Alrich, United States), and lysates were resolved on a 10% sodium dodecyl sulfate polyacrylamide gel prior to transfer onto polyvinylidene fluoride membranes. Blots were subjected to blocking in 5% milk in TBST and incubated overnight with respective primary antibodies - rabbit anti-HA (1:1500, Cat. No. H6908, Sigma-Aldrich, USA), rabbit anti-c-fos (1:1000, Cat. No. 2250, Cell Signalling Technology, USA), rabbit anti-β-actin (1:10,000, Cat. No. AC026, Abclonal Technology, USA). Blots were exposed to HRP-conjugated goat anti-rabbit secondary antibody (1:6000, Cat. No. AS014, Abclonal Technology, USA) for one hour with signal visualized on a GE Amersham Imager 600 (GE Life Sciences, USA) using a western blotting detection kit (WesternBright ECL, Advansta, USA). Densitometric quantitative analysis was performed using ImageJ software.

### Immunofluorescence analysis

Coronal brain sections (40 µm) were generated on a vibratome (Leica, Germany) from adult CamKIIα-tTA::TRE-hM4Di bigenic mice sacrificed by transcardial perfusion with 4% paraformaldehyde. Sections were permeabilized and blocked in phosphate-buffered saline with 0.3% Triton X-100 (PBSTx) containing 10% horse serum (Thermo Fisher Scientific, Cat. No. 26-050-088) for two hours at room temperature. The sections were then incubated with primary antibody for double-label immunofluorescence experiments, to examine the co-localization of the HA-tagged hM4Di-DREADD with markers for excitatory and inhibitory neurons, and glial cells in the hippocampus and neocortex. The following antibody cocktails were used: rat anti-HA (1:200, Cat. No. 10145700, Roche Diagnostics, USA) with rabbit anti-CamKIIα (1:200, Cat. No. sc-12886-R, Santa Cruz, USA), mouse anti-PV (1:500, Sigma-Aldrich, Cat. No. P3088), or rabbit anti-GFAP (1:500, Cat. No. AB5804, Chemicon, USA) for four days at 4°C. This was followed by washing of sections with 0.3% PBSTx thrice for fifteen minutes each. The sections were then incubated with the following cocktails of secondary antibodies, namely, goat anti-rat IgG conjugated to Alexa Fluor 488 (1:500; Cat. No. A-21212, Invitrogen, USA) and goat anti-rabbit IgG conjugated to Alexa Fluor 568 (1:500; Cat. No. A-11011, Invitrogen), or goat anti-rat IgG conjugated to Alexa Fluor 488 (1:500; Cat. No. A-21212, Invitrogen) and donkey anti-mouse IgG conjugated to Alexa Fluor 555 (1:500; Cat. No. A-31570, Invitrogen) for two hours at room temperature. After sequential washing with 0.3% PBSTx, sections were mounted on slides using Vectashield Antifade Mounting Medium with DAPI (H-1200, Vector Laboratories, USA) and images were visualized on a FV1200 confocal microscope (Olympus, Japan).

### Behavioral Assays

Reflex behaviors for neonates were analyzed on CamKIIα-tTA::TRE-hM4Di bigenic mouse pups commencing on P9 till P12, with air righting, surface righting and negative geotaxis determined.

#### Air righting

Animals were allowed to fall 10 times from a height of 25 cm, facing upside down. Number of correct landings, as observed by falling on all four paws, was determined. *Negative geotaxis*: Animals were placed on an inclined plank (30°), facing downwards. The amount of time taken by the animal to turn (180°) and face upwards was noted.

#### Surface righting

Time for the pup to attain a standing position with all four paws was noted when placed upside down in the home cage.

In adulthood, CamKIIα-tTA::TRE-hM4Di bigenic mice were subjected to behavioral assays to assess anxiety-like behavior (open field test - OFT; elevated plus maze test - EPM; light-dark avoidance test - LD box), despair-like behavior (tail suspension test - TST and forced swim test - FST) and sensorimotor gating behavior which was assessed via the prepulse inhibition (PPI) test. All anxiety-like behavioral assays were recorded and tracked using an overhead camera at 25 frames per second. All despair-like behaviors were recorded using a side-mounted webcam (Logitech, Switzerland). Behavior tracking was done using the automated platform Ethovision XT 11.

#### Open field test

Mice were released into one corner (chosen at random) of the open field arena (40 cm x 40 cm x 40 cm), and allowed to explore for ten minutes. The total distance moved in the arena, the percent time and percent distance in the center, and number of entries to the center were determined.

#### Elevated plus maze

Animals were introduced to the elevated plus maze for ten minutes and were placed in the center of the plus maze facing the open arms. The elevated plus maze was built such that the two arms both open and closed (30 cm x 5 cm each) were elevated 50 cm above the ground. The walls of the closed arms were 15 cm high. The total distance moved in the maze, the percent time, percent distance and number of entries in the open and closed arms were determined.

#### Light-dark avoidance test

The light-dark box was made of two joint chambers-the light chamber (25 cm x 25 cm) and the dark chamber (15 cm x 25 cm). The two areas were connected by an opening (10 cm x 10 cm). Mice were released into the arena facing the light chamber at the cusp of the lit and dark arena for a duration of ten minutes. The number of entries and the percent time spent in the light arena was assessed.

#### Tail suspension test

Animals were suspended by their tail for six minutes at a height of 50 cm above the ground, and the total immobility time and number of immobility events was assessed for a duration of five minutes excluding the first minute of the test.

#### Forced swim test

Animals were allowed to swim for six minutes in a transparent cylinder of 50 cm height and 14 cm inner diameter, filled with water (25°C) up to a height of 30 cm. The total immobility time and number of immobility events was determined for a duration of five minutes excluding the first minute of the test.

#### Prepulse inhibition test

Animals were assayed for perturbation of sensorimotor gating on the prepulse inhibition test performed using a startle and fear conditioning apparatus (Panlab, Spain). CamKIIα-tTA::TRE-hM4Di bigenic mice were allowed to habituate to the restrainer and testing apparatus for fifteen minutes daily across four days, followed by a fifteen minute habituation per day for four days with an exposure to 65 dB background white noise. On the test day, following exposure of the animals to 65 dB background white noise for five minutes, ten tone pulses (120 dB, 1 s) were presented to the mice to measure basal startle response (first block). The mice were then randomly presented with either only tone (120 dB, 1 s; x10) or a 100 ms prepulse which was either +4 dB (69dB), +8 dB (73dB) or +16 dB (81dB) higher than the background noise, five times each, which co-terminated with a 1 s, 120 dB tone. The percent PPI was calculated using the following formula: Percent PPI = 100 × (average startle response with only tone − average startle response with the prepulse) ∕ average startle response with only tone.

### Statistical analysis

All experiments had two treatment groups and were subjected to a two-tailed, unpaired Student’s *t*-test using GraphPad Prism (Graphpad Software Inc, USA). The Kolmogorov-Smirnov test was used to confirm normality of distribution. Welch correction was applied when a significant difference in the variance between groups was observed. Data are expressed as mean ± standard error of the mean (S.E.M) and statistical significance was set at *p* < 0.05.

## Results

### Selective expression of hM4Di-DREADD in CamKIIα-positive forebrain excitatory neurons in CamKIIα-tTA::TRE-hM4Di bigenic mice

To address the behavioral consequences of hM4Di-DREADD mediated inhibition of forebrain excitatory neurons in postnatal and juvenile windows of life, CamKIIα-tTA::TRE-hM4Di bigenic mice were generated. The expression of the HA-tagged hM4Di-DREADD was characterized in the hippocampus and cortex, wherein the CamKIIα-tTA driver would result in expression in forebrain excitatory neurons (Mayford et al., 1996; Wang et al., 2013), as well as within brain regions such as the periaqueductal gray (PAG) and pallidum, which are known to lack an expression of CamKIIα-positive neurons. The presence of the HA-tagged hM4Di-DREADD was confirmed in CamKIIα-positive neurons in both, the hippocampus (Fig. 1A) and the cortex (Fig. 1D) of adult CamKIIα-tTA::TRE-hM4Di bigenic mice. The HA-tagged hM4Di-DREADD was not present on the inhibitory parvalbumin (PV)-positive neurons (Fig. 1B), as well as in glial fibrillary acidic protein (GFAP) immunopositive astrocytes (Fig. 1C) in the hippocampus. We also confirmed the absence of the HA-tagged hM4Di-DREADD in select brain regions that lack CamKIIα-positive neurons, namely the PAG (Fig. 1E) and pallidum (Fig. 1F).

**Figure 1:**
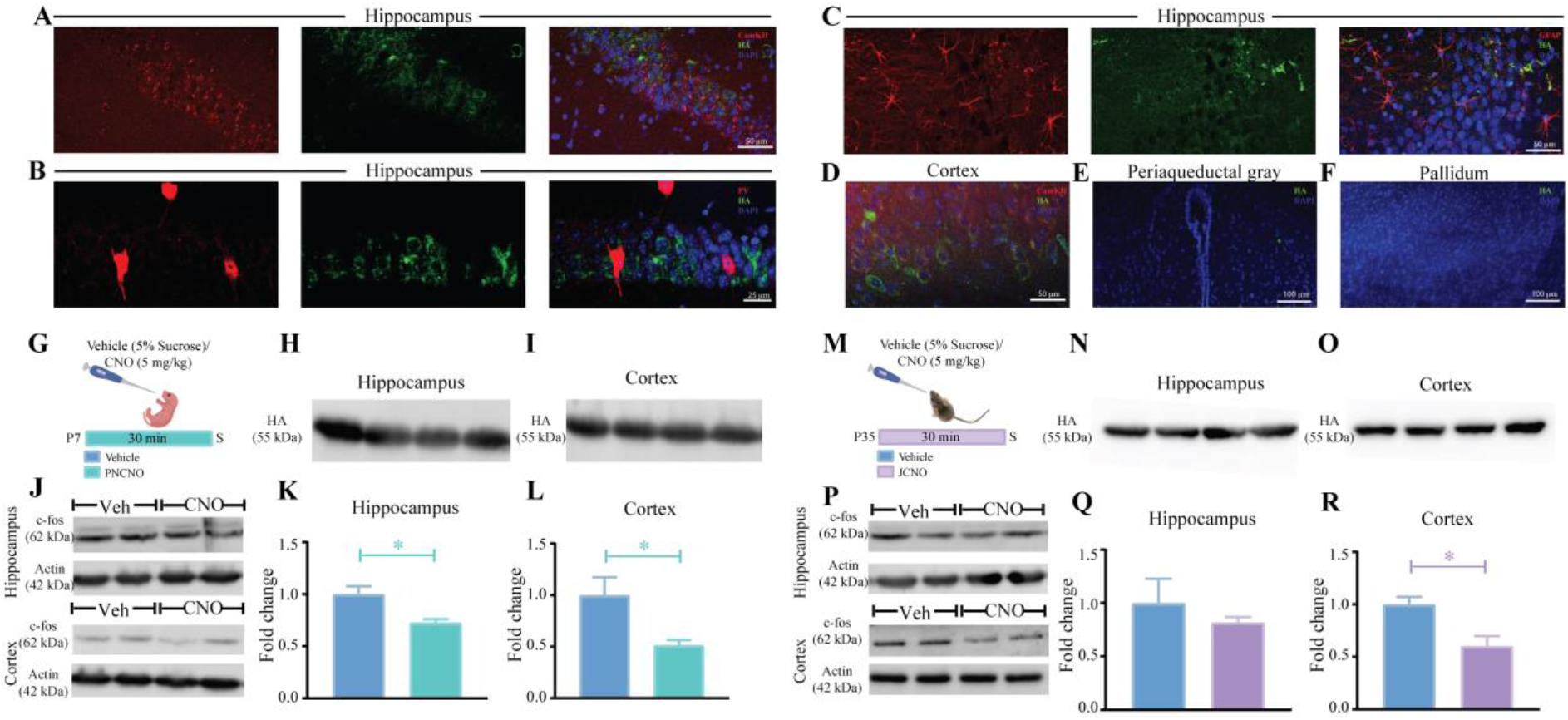
Selective expression of hM4Di-DREADD in CamKIIα-positive forebrain excitatory neurons in CamKIIα-tTA::TRE-hM4Di bigenic mice. Shown are representative confocal images indicating expression of the HA-tagged hM4Di-DREADD in the Hippocampus (A) as identified by HA/CamKIIα double immunofluorescence. HA-tagged hM4Di-DREADD expression was not observed in either Parvalbumin (PV)-positive inhibitory interneurons (B), or GFAP-positive astrocytes (C). HA-tagged hM4Di-DREADD in the cortex (D) was also observed in the CamKIIα-positive neurons as identified with HA/CamKIIα double immunofluorescence. Immunofluorescence experiments indicate the absence of expression of HA-tagged hM4Di-DREADD in subcortical brain regions, namely the periaqueductal gray (E) and pallidum (F). Shown is a schematic of the experimental paradigm for harvesting cortex and hippocampus at P7 for western blotting analysis (G). HA expression was clearly noted in the cortex (H) as well as the hippocampus (I) in western blots from CamKIIα-tTA::TRE-hM4Di bigenic pups (P7). Shown are representative western blots for c-fos along with their respective actin loading controls at P7 half an hour post vehicle (Veh) or CNO treatment for cortex (upper panel) and hippocampus (lower panel) (J). Quantitative densitometry indicated a significant reduction in c-fos protein levels in the cortex (K) as well as hippocampus (L) of PNCNO-treated bigenic pups at P7 as compared to their vehicle-treated controls. Shown is a schematic of the experimental paradigm for harvesting cortex and hippocampus at P35 in the juvenile window for western blotting analysis (M). HA expression was noted in the cortex (N) as well as the hippocampus (O) of CamKIIα-tTA::TRE-hM4Di bigenic juvenile mice (P35). Shown are representative western blots for c-fos along with their respective actin loading controls at P35 half an hour post vehicle (Veh) or CNO treatment for cortex (upper panel) and hippocampus (lower panel) (P). Quantitative densitometry indicated a significant reduction in c-fos protein levels in the cortex (Q) but not in the hippocampus (R) of JCNO-treated bigenic mice at P35 as compared with their vehicle-treated controls. All immunofluorescence experiments and western blotting experiments were performed on n = 3-5 per group. Results are expressed as the mean ± S.E.M. \**p*<0.05, as compared to vehicle-treated controls using the two-tailed, unpaired Student’s *t*-test.

We next examined the presence of the HA-tagged hM4Di-DREADD in the hippocampus and cortex using western blotting analysis. CamKIIα-tTA::TRE-hM4Di bigenic mice at postnatal day 7 (P7, Fig. 1G-I) as well as in juveniles at postnatal day 35 (P35, Fig. 1M-O) exhibited robust expression of the HA-tagged hM4Di-DREADD. We then performed western blotting analysis for the neuronal activity marker, c-fos, to examine whether hM4Di-DREADD evoked a reduction in neuronal activity in the hippocampus and cortex, half an hour post CNO or vehicle treatment to postnatal pups at P7 and juvenile animals at P35. Western blotting analysis, followed by quantitative densitometry for c-fos protein levels revealed a significant reduction in the hippocampus (Fig. 1K) and cortex (Fig. 1L) of PNCNO-treated CamKIIα-tTA::TRE-hM4Di bigenic mouse pups. For the JCNO-treated CamKIIα-tTA::TRE-hM4Di bigenic mice at P35, we did not observe a change in the c-fos protein levels in the hippocampus (Fig. 1Q), but did note a significant decline in c-fos protein levels in the cortex (Fig. 1R) of the JCNO-treated cohort. Collectively, these results indicate that the expression of HA-tagged hM4Di-DREADD is restricted to forebrain CamKIIα-positive neurons, and that treatment with the DREADD ligand CNO in the early postnatal or juvenile window evokes a decline in activity of within the forebrain regions of the hippocampus and cortex, as indicated by a reduction in protein levels of the neuronal activity marker, c-fos.

### Chronic hM4Di-DREADD mediated inhibition of CamKIIα-positive forebrain excitatory neurons during the early postnatal window does not influence anxiety-like behavior in adulthood in male or female mice

We set out to examine the behavioral influence of chronic CNO-mediated hM4Di-DREADD inhibition of CamKIIα-positive forebrain excitatory neurons during the early postnatal or juvenile window by orally administering the DREADD ligand, CNO (5 mg/kg), or vehicle to CamKIIα-tTA::TRE-hM4Di bigenic mice once daily from P2 to P14 (Fig. 2A-PNCNO), or from P28 to P40 (Fig. 2F - JCNO). PNCNO or JCNO treatments did not alter the body weight which was measured across the period of treatment (Fig. 2B, 2G). PNCNO treatment did not alter the normal ontogeny of reflex behaviors, namely air righting (Fig. 2C), negative geotaxis (Fig. 2D) and surface righting (Fig. 2E), in PNCNO-treated CamKIIα-tTA::TRE-hM4Di bigenic mouse pups as compared to their vehicle-treated controls.

**Figure 2:**
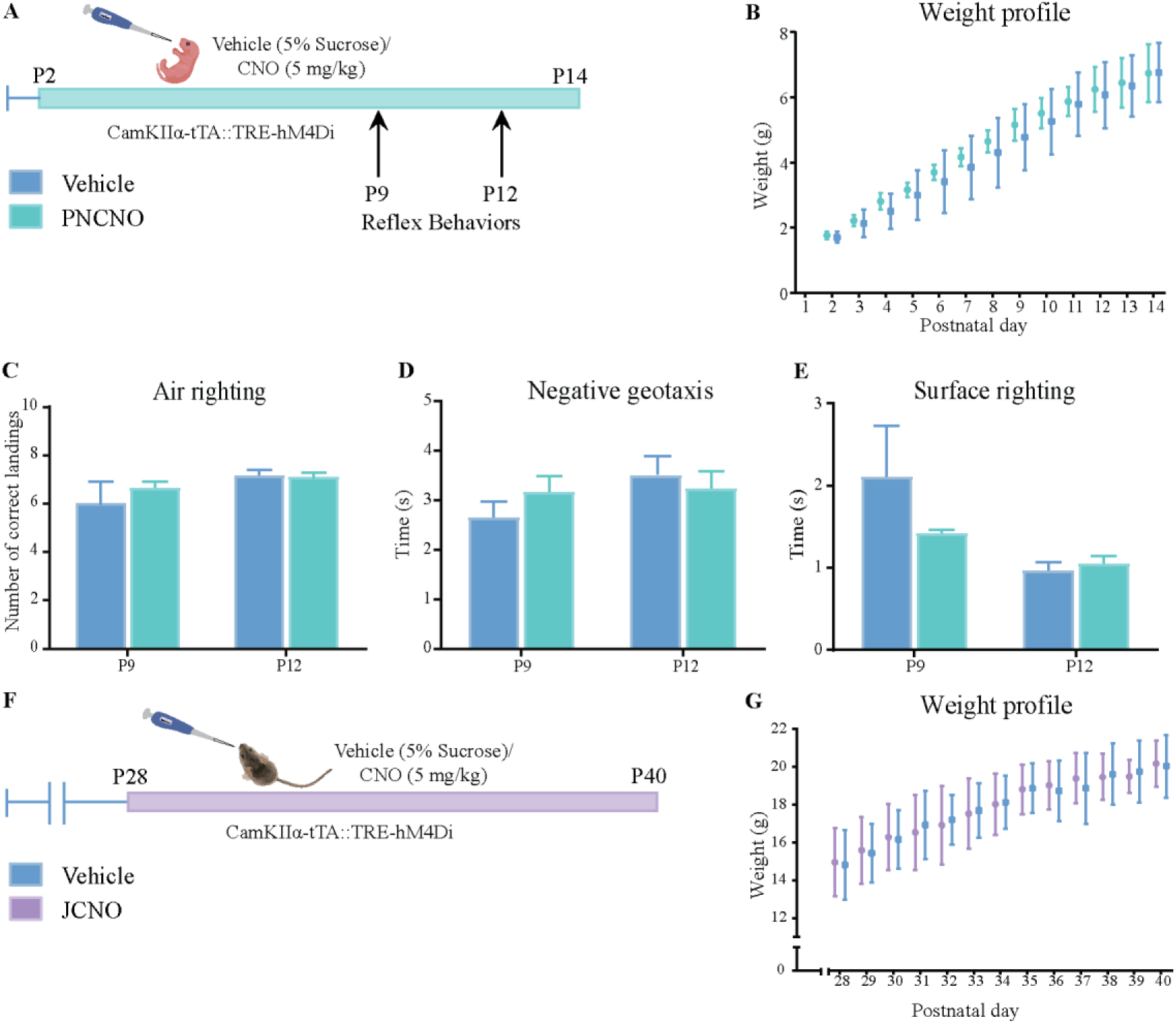
Influence of chronic hM4Di-DREADDmediated inhibition of CamKIIα-positive forebrain neurons in the early postnatal or juvenile windows on weight and reflex development. Shown is a schematic (A) for the experimental paradigm for vehicle (5% sucrose) or CNO (5 mg/kg) administration in the early postnatal window (P2-P14) in CamKIIα-tTA::TRE-hM4Di bigenic pups. Pups were assessed for weight gain across the postnatal developmental window and for reflex behaviors on postnatal days 9 and 12. No significant change was observed in the weight profile of CNO-administered pups as compared to their vehicle-treated age-matched controls across the duration of CNO treatment from postnatal day 2-14 (n = 6) (B). Reflex behaviors were not altered in PNCNO-treated CamKIIα-tTA::TRE-hM4Di bigenic pups as compared to vehicle-treated controls at P9 or P12 as assessed by determining the number of correct landings for air righting (C), and the time taken for reorientation in both negative geotaxis (D), and surface righting (E) assays. Shown is a schematic (F) for the experimental paradigm for vehicle (5% sucrose) or CNO (5 mg/kg) administration in the early juvenile window (P28-P40) to CamKIIα-tTA::TRE-hM4Di bigenic male mice. No significant change was noted in the weight profile of animals fed with CNO (5 mg/kg) once daily from P28 to P40 as compared with their vehicle-treated controls across the duration of drug treatment (n = 5-6 per group) (G). Results are expressed as mean ± S.E.M., and groups are compared using the two-tailed, unpaired Student’s *t*-test.

We next addressed whether a history of hM4Di-DREADD mediated inhibition of CamKIIα-positive forebrain excitatory neurons during the early postnatal window alters anxiety-like behavior in adulthood. We subjected CamKIIα-tTA::TRE-hM4Di bigenic adult male and female mice with a history of PNCNO treatment to a battery of behavioral tests to assess anxiety-like behavior, namely the open field test (OFT), elevated plus maze (EPM) test, and light-dark avoidance test (LD box). We noted no change in behavior in the OFT in both male (Fig. 3B-E; Ext data Fig. 3-1B) and female (Fig. 4B-E; Ext data Fig. 4-1B) CamKIIα-tTA::TRE-hM4Di bigenic mice, with no difference observed in the total distance travelled (Fig. 3C, 4C), percent time spent in the center (Fig. 3D, 4D), number of entries to the center (Fig. 3E, 4E) and percent distance travelled in the center (Ext data Fig. 3-1B, 4-1B). PNCNO-treated male and female mice also did not show any change in behavior on the EPM (Fig. 3F-J; 4F-J; Ext data Fig. 3-1C, D; Ext data Fig. 4-1C, D), with no difference observed in the percent time spent in closed arms (Fig. 3G, 4G) or open arms (Fig. 3H, 4H), as well as the percent distance travelled in open arms (Ext data Fig. 3-1C, 4-1C) or closed arms (Ext data Fig. 3-1D, 4-1D). Total number of entries to open arms (Fig. 3I, 4I) as well as closed arms (Fig. 3J, 4J) was also unaltered. Further, no difference in behavior was noted between PNCNO CamKIIα-tTA::TRE-hM4Di bigenic adult male and female mice and their respective vehicle-treated controls in the LD box in either the percent time spent in the light box (Fig. 3L, 4L), or in the number of entries to the light box (Fig. 3M, 4M). Taken together, these findings indicate that PNCNO-mediated, chronic hM4Di-DREADD inhibition of CamKIIα-positive forebrain excitatory neurons during the early postnatal window does not influence anxiety-like behavior in adulthood on the OFT, EPM and LD tests in both male and female mice.

**Figure 3:**
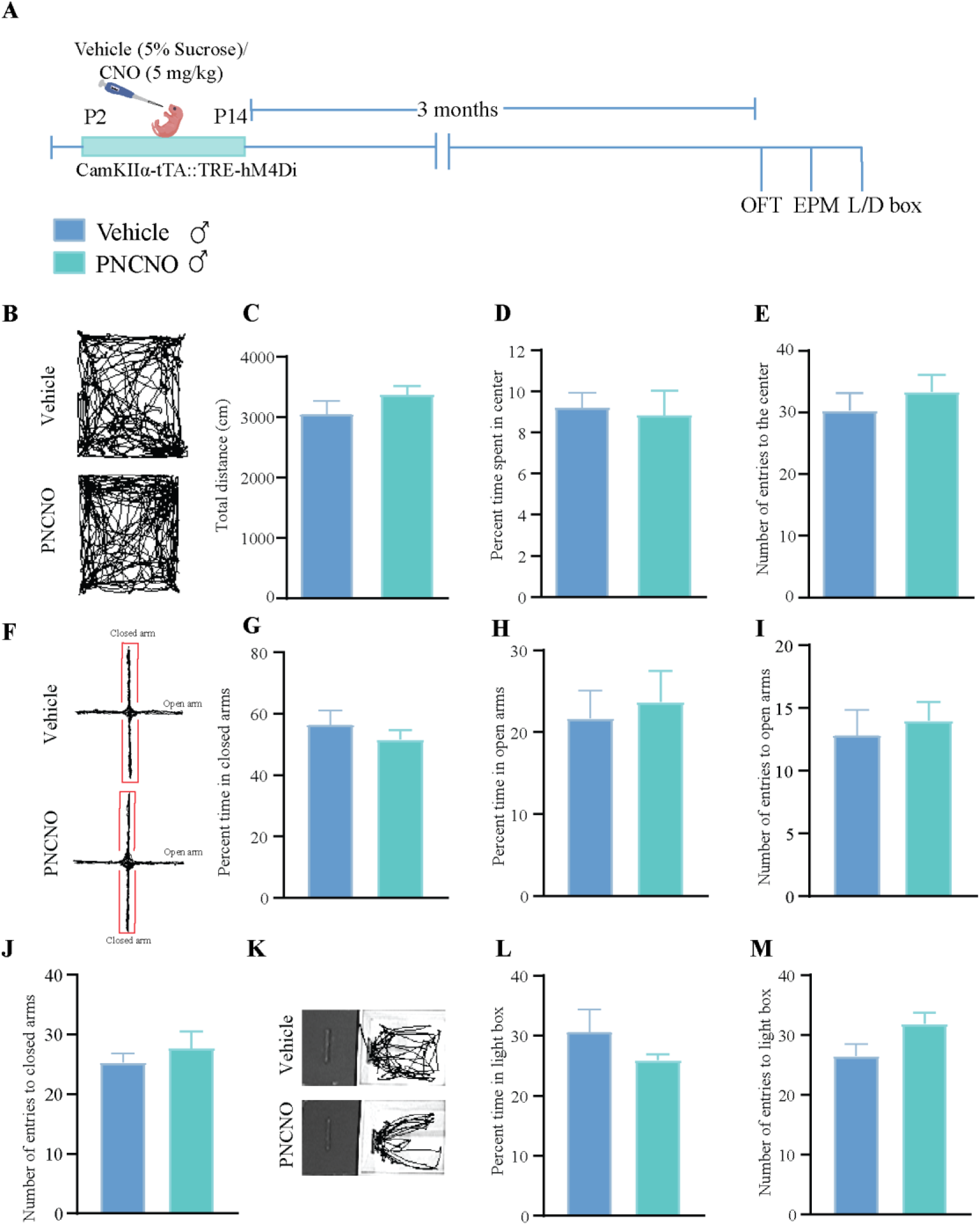
Chronic hM4Di-DREADD mediated inhibition of CamKIIα-positive forebrain excitatory neurons during the early postnatal window does not influence anxiety-like behavior in adulthood in CamKIIα-tTA::TRE-hM4Di bigenic male mice. Shown is a schematic (A) for the experimental paradigm for vehicle (5% sucrose) or CNO (5 mg/kg) administration in the early postnatal window (P2-P14) to CamKIIα-tTA::TRE-hM4Di bigenic pups which were then assessed for anxiety-like behaviors three months post cessation of CNO treatment in adulthood (A). Shown are representative tracks for vehicle-treated (top panel) and PNCNO-treated CamKIIα-tTA::TRE-hM4Di bigenic male mice (bottom panel) in the open field arena (B). No significant difference between vehicle and PNCNO-treated male mice were noted for the total distance travelled in the arena (C), percent time spent in the center (D), or total number of entries to the center of the open field arena (E) (n = 10 for vehicle and n = 12 for PNCNO male mice). Shown are representative tracks for vehicle-treated (top panel) and PNCNO-treated CamKIIα-tTA::TRE-hM4Di bigenic male mice (bottom panel) in the elevated plus maze (F). No significant difference was observed between vehicle and PNCNO-treated mice in percent time spent in the closed (G) or open (H) arms of the plus maze, as well as for the number of entries to open (I) or closed (J) arms (n = 14 for vehicle and n = 12 for PNCNO male mice). Shown are representative tracks for vehicle-treated (top panel) and PNCNO-treated CamKIIα-tTA::TRE-hM4Di bigenic male mice (bottom panel) in the light-dark box (K). No significant difference was noted between vehicle and PNCNO-treated mice in either total time spent (L), or total number of entries (M) into the light chamber of the light-dark box (n = 14 for vehicle and n = 12 for PNCNO male mice). Results are expressed as mean ± S.E.M., and groups are compared using the two-tailed, unpaired Student’s *t*-test.

**Figure 4:**
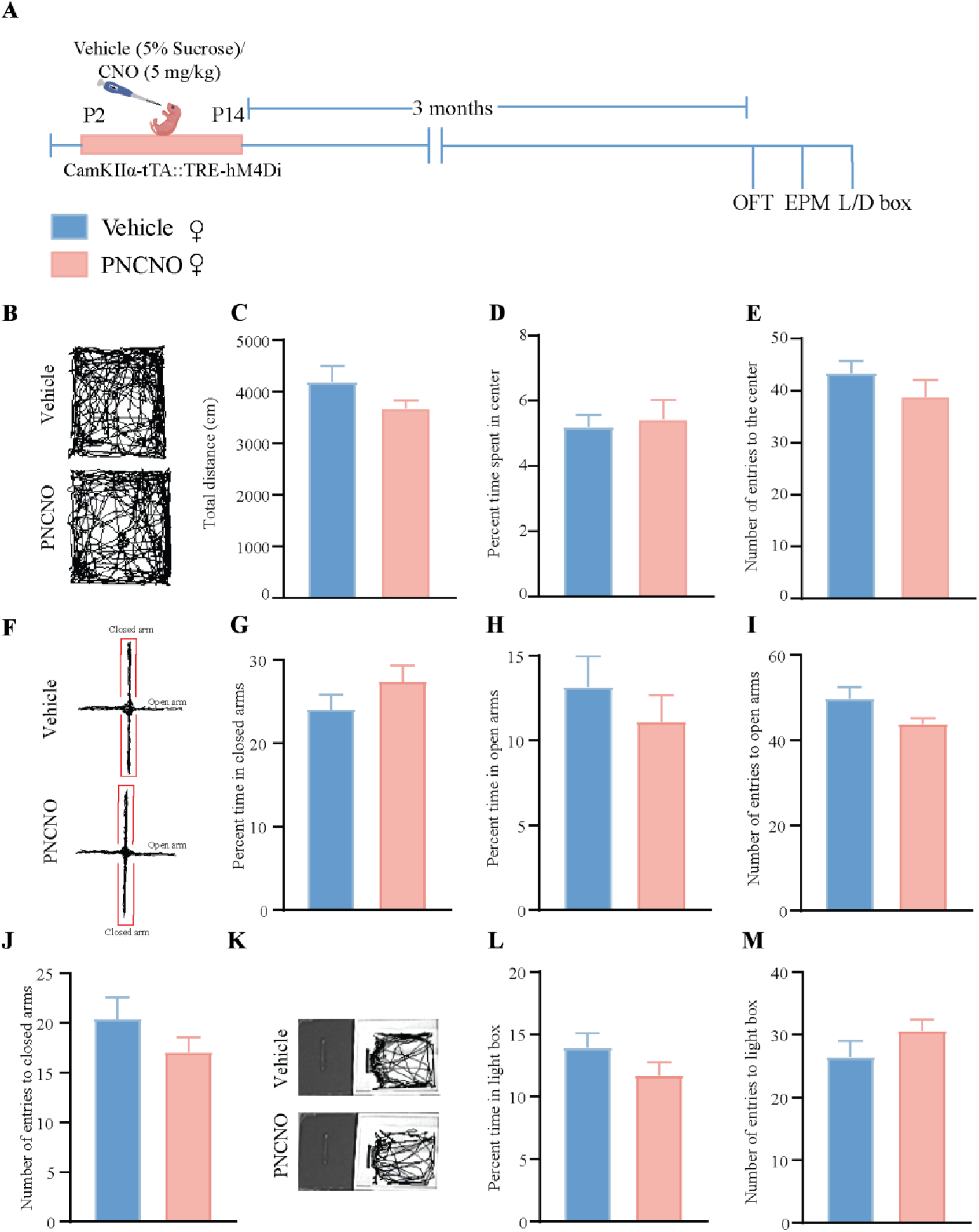
Chronic hM4Di-DREADD mediated inhibition of CamKIIα-positive forebrain excitatory neurons during the early postnatal window does not influence anxiety-like behavior in adulthood in CamKIIα-tTA::TRE-hM4Di bigenic female mice. Shown is a schematic (A) for the experimental paradigm for vehicle (5% sucrose) or CNO (5mg/kg) administration in the early postnatal window (P2-P14) in CamKIIα-tTA::TRE-hM4Di bigenic pups which were then assessed for anxiety-like behaviors three months post cessation of CNO treatment in adulthood in the female cohort (A). Shown are representative tracks for vehicle-treated (top panel) and PNCNO-treated CamKIIα-tTA::TRE-hM4Di bigenic female mice (bottom panel) in the open field arena (B). No significant difference between vehicle and PNCNO-treated female mice was noted for the total distance travelled in the arena (C), percent time spent in the center (D), or total number of entries to the center of the open field arena (E) (n = 10 for vehicle and n = 12 for PNCNO female mice). Shown are representative tracks for vehicle-treated (top panel) and PNCNO-treated CamKIIα-tTA::TRE-hM4Di bigenic female mice (bottom panel) in the elevated plus maze (F). No significant difference was observed between vehicle and PNCNO-treated female mice in the percent time spent in the closed (G) or open (H) arms of the plus maze, as well as the number of entries to open (I) or closed (J) arms (n = 12 for vehicle and n = 11 for PNCNO female mice). Shown are representative tracks for vehicle-treated (top panel) and PNCNO-treated CamKIIα-tTA::TRE-hM4Di bigenic female mice (bottom panel) in the light-dark box (K). No significant difference was noted between vehicle and PNCNO-treated female mice in either total time spent (L), or total number of entries (M) into the light chamber of the light-dark box (n = 14 for vehicle and n = 12 for PNCNO female mice). Results are expressed as mean ± S.E.M., and groups are compared using the two-tailed, unpaired Student’s *t*-test.

### Chronic hM4Di-DREADD mediated inhibition of CamKIIα-positive forebrain excitatory neurons during the juvenile window does not influence anxiety-like behavior in adulthood

We examined whether chronic CNO-mediated hM4Di-DREADD inhibition of CamKIIα-positive forebrain excitatory neurons during the juvenile window (P28-P40) alters anxiety-like behavior in adulthood. We subjected CamKIIα-tTA::TRE-hM4Di bigenic adult male mice with a history of JCNO treatment to a battery of behavioral tests commencing two months after the cessation of CNO treatment. We examined anxiety-like behavior on the OFT, EPM and LD tests. JCNO-treated CamKIIα-tTA::TRE-hM4Di bigenic adult male mice did not exhibit any difference in anxiety-like behavior on these behavioral tasks (Fig. 5A). On the OFT, we noted no change in the total distance travelled in the OFT arena (Fig. 5B, C), as well as in the percent time spent in the center (Fig. 5D), the percent distance travelled in the center (Ext data Fig. 5-1B), or in the total number of entries to the center of the OFT arena (Fig. 5E). We also observed no significant differences in behavior on the EPM, with no change noted for the percent time spent in closed arms (Fig. 5G) or open arms (Fig. 5H), as well as the percent distance travelled in open arms (Ext data Fig. 5-1C) or closed arms (Ext data Fig. 5-1D). The total number of entries to both the open (Fig. 5I) and closed arms (Fig. 5J) was also unchanged across vehicles and JCNO-treated CamKIIα-tTA::TRE-hM4Di bigenic adult male mice. Behavioral measures assessed on the LD box were also not altered between vehicle and JCNO-treated CamKIIα-tTA::TRE-hM4Di bigenic adult male cohorts, with no difference noted for either the percent time spent in the light box (Fig. 5L), or the number of entries to the light box (Fig. 5M).

**Figure 5:**
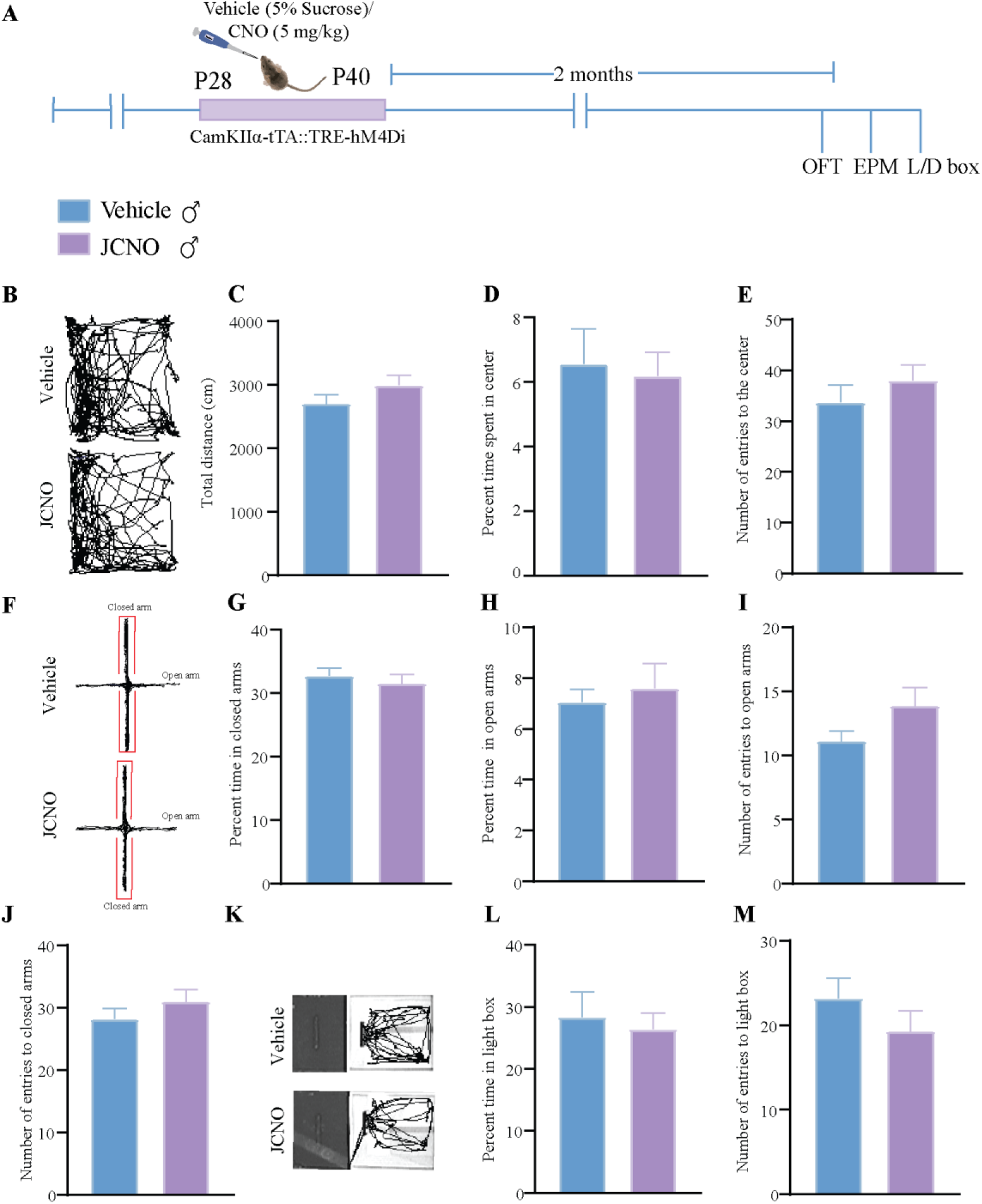
Chronic hM4Di-DREADD mediated inhibition of CamKIIα-positive forebrain excitatory neurons during the juvenile window does not influence anxiety-like behavior in adulthood in CamKIIα-tTA::TRE-hM4Di bigenic male mice. Shown is a schematic (A) for the experimental paradigm for vehicle (5% sucrose) or CNO (5 mg/kg) administration in the juvenile window (P28-P40) to CamKIIα-tTA::TRE-hM4Di bigenic mice, which were then assessed for anxiety-like behaviors two months post cessation of CNO treatment in adulthood in the male cohort (A). Shown are representative tracks for vehicle-treated (top panel) and JCNO-treated CamKIIα-tTA::TRE-hM4Di bigenic male mice (bottom panel) in the open field arena (B). No significant difference was noted between vehicle and JCNO-treated male mice in the total distance travelled in the arena (C), percent time spent in the center (D), or total number of entries to the center of the open field arena (E) (n = 18 for vehicle and n = 16 for JCNO male mice). Shown are representative tracks for vehicle-treated (top panel) and JCNO-treated CamKIIα-tTA::TRE-hM4Di bigenic male mice (bottom panel) in the elevated plus maze (F). No significant difference was observed between vehicle and JCNO-treated male mice in percent time spent in the closed (G) or open (H) arms of the plus maze, as well as for the number of entries to open (I) or closed (J) arms (n = 18 for vehicle and n = 16 for JCNO male mice). Shown are representative tracks for vehicle-treated (top panel) and JCNO-treated CamKIIα-tTA::TRE-hM4Di bigenic male mice (bottom panel) in the light-dark box (K). No significant difference was noted between vehicle and JCNO-treated male mice in either total time spent (L), or total number of entries (M) into the light chamber of the light-dark box (n = 18 for vehicle and n = 16 for JCNO male mice). Results are expressed as mean ± S.E.M., and groups are compared using the two-tailed, unpaired Student’s *t*-test.

### Chronic hM4Di-DREADD mediated inhibition of CamKIIα-positive forebrain excitatory neurons during the early postnatal or juvenile window does not influence despair-like behavior in adult mice

We next addressed whether a history of hM4Di-DREADD mediated inhibition of CamKIIα-positive forebrain excitatory neurons during the early postnatal or juvenile window alters despair-like behavior in adulthood on both the forced swim (FST) and the tail suspension test (TST). CamKIIα-tTA::TRE-hM4Di bigenic adult male and female mice with a history of PNCNO treatment did not show any change in either the percent immobility time (Fig. 6B, E) or the number of immobility events (Fig. 6C, F) on the FST. We also observed no change in the percent immobility time on the TST between PNCNO-treated CamKIIα-tTA::TRE-hM4Di bigenic adult male as compared to their vehicle-treated controls (Percent immobility time: Vehicle-treated CamKIIα-tTA::TRE-hM4Di bigenic adult male mice = 60.54 ± 4.5, n = 8; PNCNO-treated CamKIIα-tTA::TRE-hM4Di bigenic adult male mice = 62.56 ± 2.63, n = 9; results are expressed as mean ± S.E.M). Further, we examined whether CamKIIα-tTA::TRE-hM4Di bigenic adult male mice with a history of JCNO treatment differed from their vehicle-treated controls on the FST and TST. JCNO-treated CamKIIα-tTA::TRE-hM4Di bigenic adult male did not show any significant differences in the percent immobility time (Fig. 6H) or the number of immobility events (Fig. 6I) on the FST. We also noted no significant differences in the percent immobility time on the TST between JCNO-treated CamKIIα-tTA::TRE-hM4Di bigenic adult male and the vehicle-treated cohort (Percent immobility time: Vehicle-treated CamKIIα-tTA::TRE-hM4Di bigenic adult male mice = 64.15 ± 5.56, n = 9; JCNO-treated CamKIIα-tTA::TRE-hM4Di bigenic adult male mice = 60.52 ± 2.32, n = 9; results are expressed as mean ± S.E.M). Collectively, these results indicate that chronic hM4Di-DREADD mediated chemogenetic inhibition of CamKIIα-positive forebrain excitatory neurons during either the early postnatal or juvenile window does not alter despair-like behavior on the FST or TST in adulthood.

**Figure 6:**
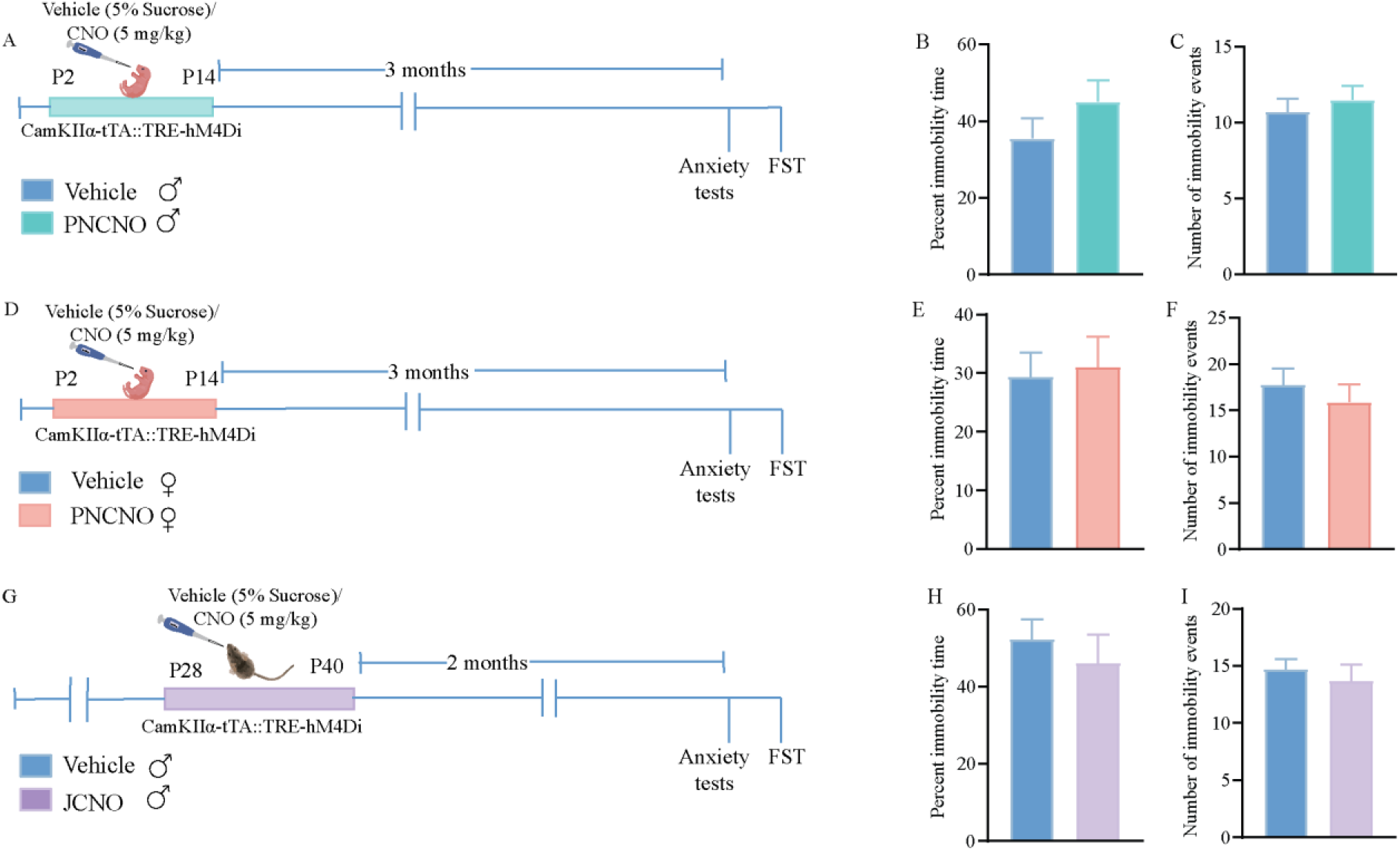
Chronic chemogenetic inhibition of CamKIIα-positive forebrain excitatory neurons during the early postnatal window as well as juvenile window does not influence despair-like behavior in adult mice. Shown is a schematic for the experimental paradigm for vehicle (5% sucrose) or CNO (5 mg/kg) administration in the early postnatal window (P2-P14) to CamKIIα-tTA::TRE-hM4Di bigenic male pups (A) which were then assessed for despair-like behavior in adulthood using the forced swim test (FST). No significant difference was noted between vehicle and PNCNO-treated male mice for either the percent time spent immobile (B), or the total number of immobility events (C) (n = 14 for vehicle and n = 12 for PNCNO male mice). Shown is a schematic for the experimental paradigm for vehicle (5% sucrose) or CNO (5 mg/kg) administration in the early postnatal window (P2-P14) in CamKIIα-tTA::TRE-hM4Di bigenic female pups (D). No significant difference was noted between vehicle and PNCNO-treated female mice for the percent time spent immobile (E), or the total number of immobility events (F) (n = 10 for vehicle and n = 10 for PNCNO female mice). Shown is a schematic for the experimental paradigm for vehicle (5% sucrose) or CNO (5 mg/kg) administration in the juvenile window (P28-P40) in CamKIIα-tTA::TRE-hM4Di bigenic male mice (G) which were then assessed for despair-like behavior in adulthood using the FST. No significant difference was noted between vehicle and PNCNO-treated male mice for the percent time spent immobile (H), or the total number of immobility events (I) (n = 11 for vehicle and n = 11 for JCNO male mice). Results are expressed as mean ± S.E.M., and groups are compared using the two-tailed, unpaired Student’s *t*-test.

### Chronic hM4Di-DREADD mediated inhibition of CamKIIα-positive forebrain excitatory neurons during the postnatal or juvenile window does not influence sensorimotor gating in adulthood in CamKIIα-tTA::TRE-hM4Di bigenic male mice

Disruption of excitation/inhibition balance in the neocortex has been linked to altered schizoaffective behavior in adulthood (Anticevic and Lisman, 2017; Fenton, 2015; Marín, 2016; Pati et al., 2020; Yizhar et al., 2011). We sought to address whether chronic CNO-mediated hM4Di inhibition of CamKIIα-positive forebrain excitatory neurons in the early postnatal or in the juvenile window resulted in any change in sensorimotor gating behavior in adulthood. To measure changes in sensorimotor gating we subjected PNCNO and JCNO-treated CamKIIα-tTA::TRE-hM4Di bigenic male mice and their respective vehicle-treated control groups to the prepulse inhibition (PPI) paradigm (Fig. 7A). We noted no significant difference in percent prepulse inhibition, at all prepulse tones tested above the background noise, between the PNCNO-treated and JCNO-treated CamKIIα-tTA::TRE-hM4Di bigenic male mice and their respective vehicle-treated controls (Fig. 7D, G). While we noted no significant alterations in the basal startle response across treatment groups in the JCNO experiment (Fig. 7F), we did observe a significant increase in basal startle response in the CamKIIα-tTA::TRE-hM4Di bigenic male mice with a history of PNCNO treatment when compared to their vehicle-treated control (Fig. 7C). These findings indicate that chronic CNO-mediated hM4Di-DREADD inhibition of CamKIIα-positive forebrain excitatory neurons during the early postnatal or juvenile window results does not result in any significant change in sensorimotor gating, but may evoke perturbed baseline startle responses in the PNCNO-treated group.

**Figure 7:**
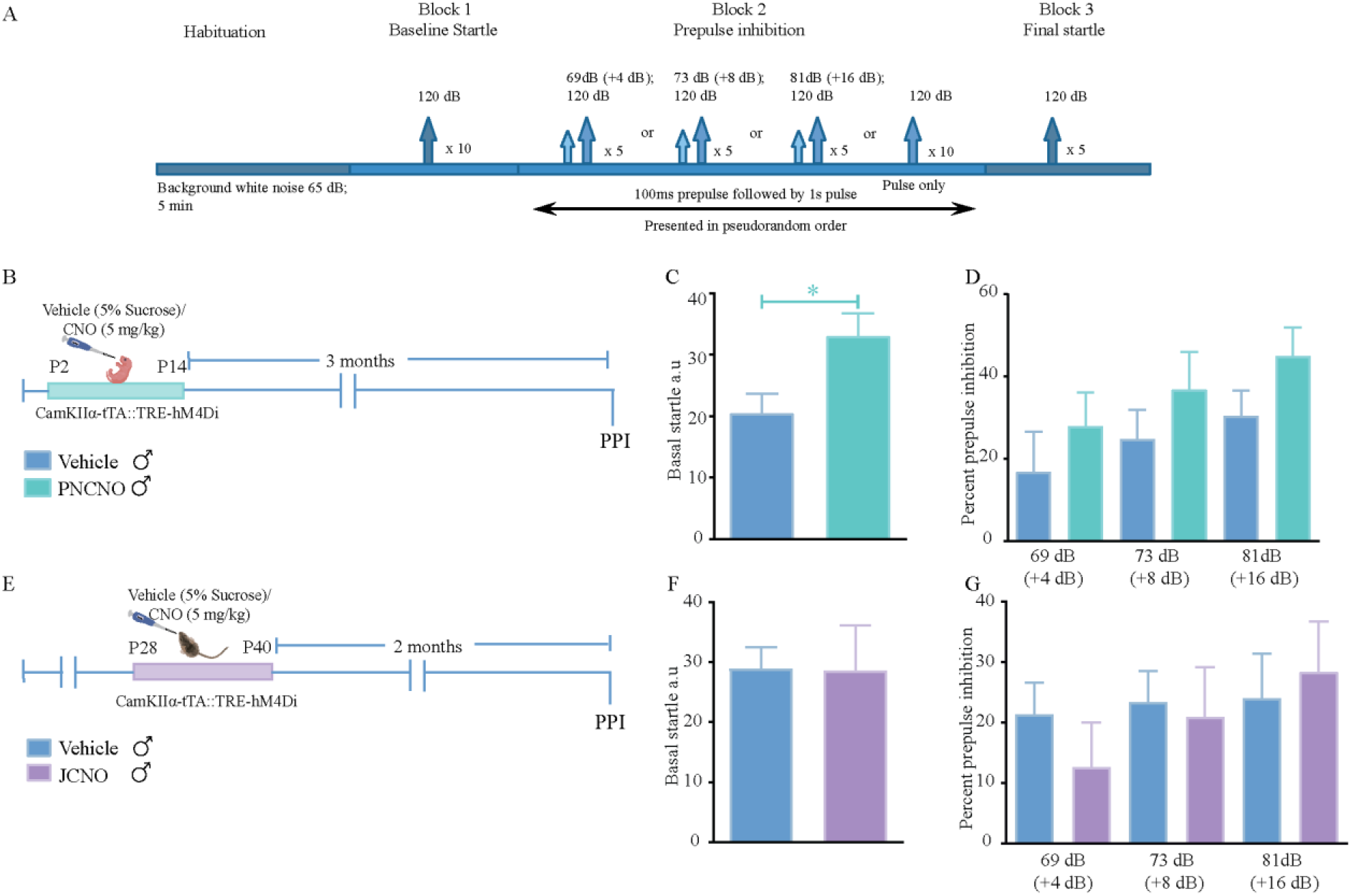
Chronic hM4Di-DREADD mediated inhibition of CamKIIα-positive forebrain excitatory neurons during the postnatal or juvenile window does not influence prepulse inhibition in adulthood in CamKIIα-tTA::TRE-hM4Di bigenic male mice. Shown is a schematic for the protocol used for prepulse Inhibition (PPI) to assess sensorimotor gating responses in adult male mice (A). PPI testing was carried out as described in Materials and Methods with basal startle determined following habituation, and PPI determined for + 4 dB (69 dB), + 8 dB (73 dB), and + 16 dB (81 dB) above background noise (65 dB), followed by exposure to 120 dB for final startle. Shown is a schematic for the experimental paradigm for administration of CNO (5 mg/kg) or vehicle (5% sucrose) administration in early postnatal window (P2-P14) to CamKIIα-tTA::TRE-hM4Di bigenic mouse pups, with PPI testing carried out four months later in adulthood in CamKIIα-tTA::TRE-hM4Di bigenic male mice (B). PNCNO-treated adult CamKIIα-tTA::TRE-hM4Di bigenic male mice show a significant increase in basal startle response (C) as compared to vehicle-treated controls. No significant differences were noted in sensorimotor gating between vehicle and PNCNO-treated CamKIIα-tTA::TRE-hM4Di bigenic male mice (D) (n = 10 per group). Shown is a schematic for the experimental paradigm for CNO (5 mg/kg) or vehicle (5% sucrose) administration in the juvenile window (P28-P40) to CamKIIα-tTA::TRE-hM4Di bigenic male mice that were subjected to PPI testing three months later (E). Basal startle response in JCNO-treated CamKIIα-tTA::TRE-hM4Di bigenic male mice was unaltered as compared to vehicle-treated CamKIIα-tTA::TRE-hM4Di bigenic male controls (F). CamKIIα-tTA::TRE-hM4Di bigenic male mice with a history of JCNO treatment did not show any change in PPI as compared to the vehicle-treated controls (G) (n = 10 for vehicle and n = 9 for JCNO male mice). Results are expressed as the mean ± S.E.M. \**p*<0.05, as compared to vehicle-treated controls using the two-tailed, unpaired Student’s *t*-test.

## Discussion

Our findings indicate that chronic hM4Di-DREADD mediated inhibition of CamKIIα-positive forebrain excitatory neurons during the early postnatal or juvenile windows of life does not influence the development of anxiety or despair-like behavior, or alter sensorimotor gating in adulthood. Preclinical studies using rodent models indicate that the first two weeks of life represent a critical period window (Baram et al., 1997), wherein the early stress of maternal separation (MS) (Benekareddy et al., 2011; De Melo et al., 2018; Kalinichev et al., 2002; Teissier et al., 2015) or pharmacological perturbations that elevate serotonin, such as postnatal selective serotonin reuptake inhibitor (SSRI) administration (Ansorge et al., 2004; Ko et al., 2014; Rebello et al., 2014; Sarkar et al., 2014; Soiza-Reilly et al., 2018), can result in the life-long programming of persistent mood-related behavioral changes. Converging evidence across diverse models of early stress has implicated perturbations in GPCR signaling during these critical periods in the establishment and eventual emergence of disrupted anxio-depressive behaviors (Benekareddy et al., 2011; Pati et al., 2020; Sarkar et al., 2014; Soiza-Reilly et al., 2018; Teissier et al., 2019; Vinkers et al., 2010a). This has led to a hypothesis that a balance between Gq and Gi-mediated GPCR signaling within neocortical brain regions during these early developmental windows may be important to shaping the development of trait anxiety and behavioral despair (Lambe et al., 2011; Tiwari et al., 2021). A recent study has shown that enhanced Gq-signaling via chemogenetic hM3Dq-DREADD mediated activation of CamKIIα-positive forebrain excitatory neurons during the postnatal, but not the juvenile or adult, temporal windows results in long-lasting increases in anxiety and despair-like behavior, accompanied by perturbed sensorimotor gating and pre-pulse inhibition (PPI) deficits (Pati et al., 2020). While several studies have used pharmacological or genetic perturbation studies to examine the contribution of Gi-coupled GPCRs, in particular the 5-HT_1A_ receptor (Garcia-Garcia et al., 2014; Gross et al., 2002; Richardson-Jones et al., 2011, 2010; Sarkar et al., 2014; Vinkers et al., 2010a), during postnatal life in programming mood-related behavior, this has not been directly tested using a chemogenetic based approach to perturb Gi-signaling in CamKIIα-positive forebrain excitatory neurons. Based on prior evidence that pharmacological blockade (Sarkar et al., 2014; Vinkers et al., 2010a) or genetic loss of function of the Gi-coupled 5-HT_1A_ receptor results in enhanced anxiety and despair-like behavior (Gross et al., 2002; Richardson-Jones et al., 2011, 2010), a working hypothesis would suggest the possibility that enhancing Gi-signaling in CamKIIα-positive forebrain excitatory neurons during the postnatal window might evoke a decline in anxiety and despair-like behaviors in adulthood. Prior evidence indicates that transient hM4Di-DREADD inhibition of the amygdala in infant rhesus monkeys has long-lasting effects on emotionality, with a decline noted in fear and anxiety responses (Raper et al., 2019). A study also shows that constitutive overexpression of the Gi-coupled 5-HT_1A_ receptors (Kusserow et al., 2004) can program decreased anxiety-like behavior in adulthood. This differs from the pharmacological studies, wherein 5-HT_1A_ receptor stimulation in the postnatal window using the agonist 8-OH-DPAT alone does not modulate anxiety-like behavior, but can increase despair-like behavior in adulthood (Ishikawa and Shiga, 2017), whereas blockade of 5-HT_1A_ receptors in early postnatal life with the selective antagonist, WAY 100635 evokes increased anxiety-like behavior (Sarkar et al., 2014; Vinkers et al., 2010a). However, there is a paucity of literature that addresses directly whether broad alteration of Gi-signaling within forebrain neurocircuits during the postnatal temporal window contributes to the programming of altered mood-related behaviors. The results of the present study clearly demonstrate that Gi-mediated inhibition of forebrain excitatory neurons using the hM4Di-DREADD during either the postnatal (P2-P14) or juvenile (P28-P40) windows results in no behavioral change on conflict-based tasks assessing anxiety-like behavior, namely the OFT, EPM and LD avoidance test in adulthood. Further, we also noted no change in despair-like behavior on the TST and FST, or in sensorimotor gating behavior on the PPI in adulthood.

The expression of the hM4Di-DREADD was restricted to the neocortex and hippocampus as indicated by both western-blotting and immunofluorescence analysis, and the hM4Di-DREADD expression was restricted to CamKIIα-positive forebrain excitatory neurons, and not observed in PV-positive inhibitory interneurons or GFAP-positive astrocytes (Fig. 1). Western blotting analysis for the neuronal activity marker c-fos indicates that as reported previously (Salvi et al., 2019; Vetere et al., 2017), treatment with the DREADD ligand, CNO results in reduced neuronal activity in the forebrain of CamKIIα-tTA::TRE-hM4D as indicated by a decline in c-fos protein levels. Further, we found that administration of the exogenous DREADD ligand, CNO, in the postnatal or juvenile window did not appear to alter the growth and development of animals, based on observations of no weight change in animals, as well as a normal ontogenic development of reflex behaviors in CNO-treated CamKIIα-tTA::TRE-hM4Di bigenic rat pups. This suggests that enhanced hM4Di-DREADD mediated inhibition of CamKIIα-positive forebrain excitatory neurons in postnatal life does not appear to influence the emergence of critical reflexes such as air-righting, negative geotaxis and surface righting. This is in agreement with prior studies that indicate that the DREADD agonist CNO during the postnatal window does not appear to influence the emergence of key developmental milestones (Pati et al., 2020).

A change in excitation-inhibition (E/I) balance during critical developmental time windows, with a shift towards enhanced excitation of forebrain pyramidal neurons and a commensurate reduction in inhibitory tone, has been posited to play a crucial role in the programming of life-long perturbations of mood-related behaviors in several neurodevelopmental disorder models (Fenton, 2015; Nelson and Valakh, 2015; Sohal and Rubenstein, 2019; Tatti et al., 2017; Yizhar et al., 2011). Indeed, hyperexcitation of Emx1-positive neurons from P4-14 in the neocortex using either a non-invasive bioluminescent chemogenetics approach (Medendorp et al., 2021) or hM3Dq-DREADD mediated activation of CamKIIα-positive forebrain excitatory neurons from P2-P14 (Pati et al., 2020) resulted in enhanced anxiety-like behaviors and perturbed social behavior. An important experimental counterpart would be to increase inhibition in forebrain pyramidal neurons in these developmental windows, and address the influence on the emergence of mood-related behaviors. Here we provide evidence of the consequences of enhanced hM4Di-mediated DREADD inhibition of forebrain excitatory neurons on the emergence of mood-related behavior, and indicate that this perturbation does not appear to influence the development of either anxiety or despair-like behaviors. A prior study using viral-based targeting strategies to evoke hM4Di-mediated DREADD inhibition of neurons within the medial prefrontal cortex (mPFC) during the postnatal developmental window indicated that this perturbation enhanced anxiety and despair-like behavior in adulthood, whereas hM3Dq-mediated DREADD activation overlapping with the stress of MS, ameliorated the behavioral consequences of early stress (Teissier et al., 2019). Further, a recent report also examined the influence of chemogenetic inhibition of PFC neurons, that are transiently positive for the serotonin transporter in postnatal life, and showed that the enhanced anxio-depressive behaviors noted in adult animals with a history of postnatal SSRI exposure was exacerbated upon hM4Di-DREADD mediated inhibition of this subclass of PFC neurons in adulthood. In contrast, hM3Dq-DREADD mediated activation of these PFC neurons in adulthood ameliorated the postnatal SSRI-evoked anxiety and despair-like behaviors (Soiza-Reilly et al., 2018). It is of importance to note that the promoters used in these studies in specific cases target both excitatory and inhibitory neurons of the PFC or a subclass of raphe-projecting PFC neurons, further the use of transgenic mice versus viral-mediated gene delivery, and differences in the developmental epoch targeted and nature of experimental paradigms makes it challenging to directly compare our findings with these studies, in particular given that we targeted hM4Di-mediated DREADD expression to all CamKIIα-positive forebrain excitatory neurons and these prior studies assessed effects on a subset of PFC neurons.

While we observed no change in anxiety-like behavior on the OFT, EPM and LD avoidance test, we did note that PNCNO-treated CamKIIα-tTA::TRE-hM4Di bigenic male mice exhibited a higher basal acoustic startle response, although they showed no change in PPI behavior. An enhanced baseline acoustic startle response has been suggested to be reflective of enhanced anxiety-like behavior (Grillon, 2008), however we see no indication of perturbation in anxiety on any of the conflict-anxiety behavioral tasks. We cannot preclude the possibility of a developmental perturbation of acoustic sensory circuits, given the driver we have used is broad-based, and the hM4Di-DREADD would be driven in all forebrain excitatory neurons. Collectively, our results suggest that enhancing Gi signaling in forebrain excitatory neurons during the postnatal window does not influence the programming of mood-related behaviors, however it is vital to keep in mind that this is not the same as driving enhanced Gi signaling via a specific GPCR, such as the 5-HT_1A_ receptor, and indeed it is possible that a more targeted approach to selectively enhance 5-HT_1A_ receptor signaling in forebrain excitatory neurons in this developmental window could exert a role in programming changes in emotionality.

In our study we have also addressed whether hM4Di-DREADD mediated inhibition of forebrain excitatory neurons in another critical temporal window implicated in shaping mood-related behavioral traits, namely the juvenile window, could impact anxiety and despair-like behaviors (Albrecht et al., 2017; Brydges et al., 2012, 2014; Hollis et al., 2013; Luo et al., 2014; Suri et al., 2014). Animals subjected to stress during the juvenile window exhibit increased anxiety-like behavior, show enhanced benzodiazepine sensitivity, and can establish a heightened vulnerability to adult-onset stress (Avital and Richter-Levin, 2004). Peripubertal stress also causes alterations in GABAergic neurotransmission (Tzanoulinou et al., 2014), and GAD65 haplodeficiency has been shown to be associated with resilience to juvenile stress-induced increased anxiety-like behavior, possibily due to delayed maturation of inhibitory signaling (Müller et al., 2014). Previous studies have shown that enhanced Gq-mediated signaling via a chemogenetic approach in the juvenile window does not influence anxiety, despair, or schizophrenia-like behavior (Pati et al., 2020). The present work indicated that hM4Di-DREADD inhibition of forebrain excitatory neurons during the juvenile temporal window does not appear to alter anxiety or despair-like behaviors, and does not influence sensorimotor gating. We have carried out extensive profiling of the behavioral consequences of hM4Di-DREADD inhibition of forebrain excitatory neurons in both postnatal or juvenile life, and the results demonstrate that this perturbation does not alter the emergence of anxiety, despair and schizophrenia-like behavior on a variety of behavioral tasks in adulthood. This differs quite starkly from our recent study wherein hM3Dq-DREADD mediated activation of forebrain excitatory neurons in postnatal, but not juvenile or adult, life resulted in persistent increases in anxiety, despair and schizophrenia-like behavior, accompanied by specific molecular, metabolic and functional changes in the both the neocortex and the hippocampus (Pati et al., 2020). While we do not observe any change in mood-related behaviors following hM4Di-DREADD inhibition of forebrain excitatory neurons, we cannot preclude the possibility that these animals may exhibit differential responses to a stressor experience in adulthood. Our work motivates future investigation to address in detail how perturbations in GPCR signaling within forebrain circuits during critical developmental time-windows may shape vulnerability or resilience to adult-onset stressors.

---

**Extended data Figure 3-1:**
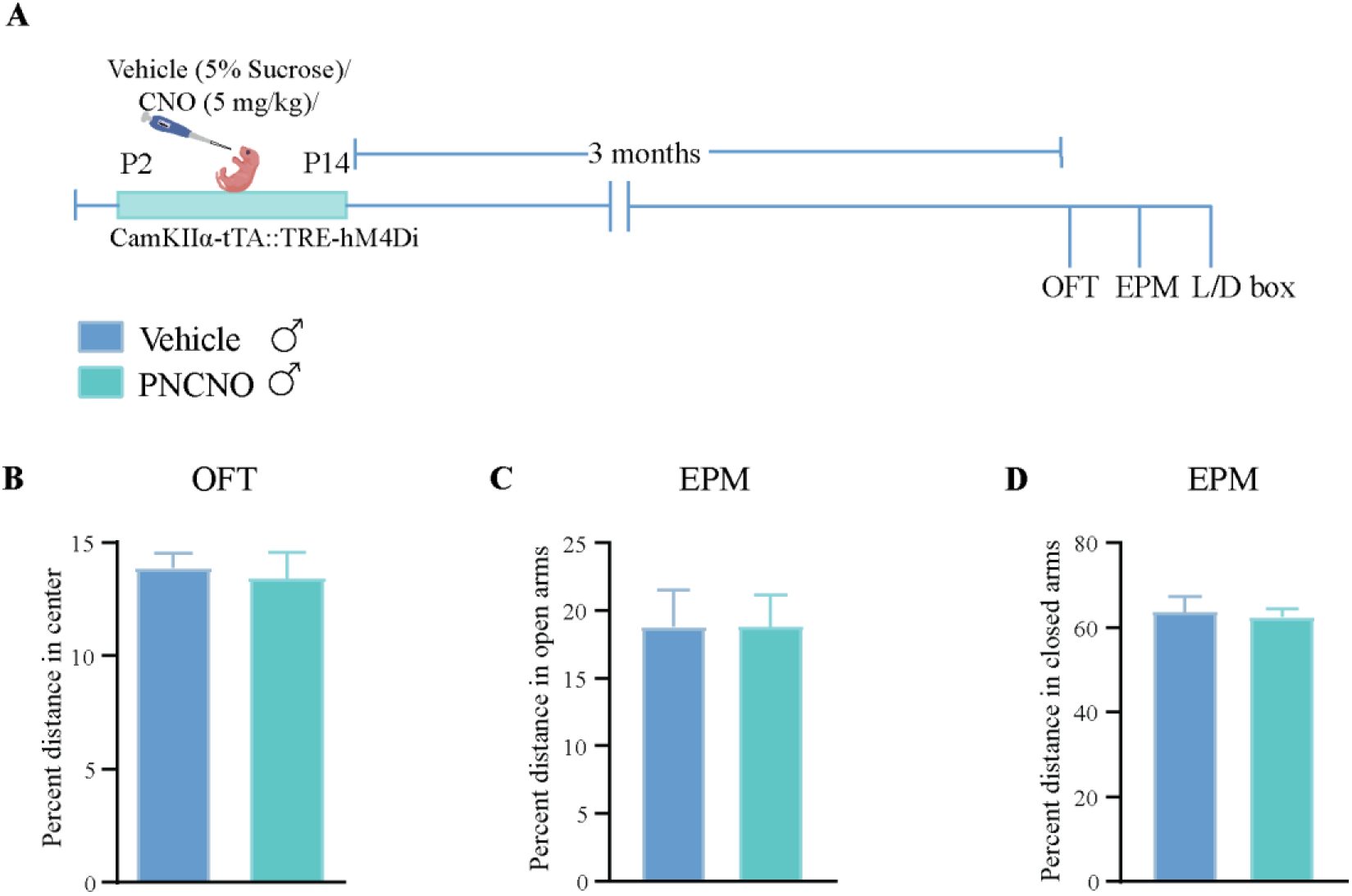
Chronic hM4Di-DREADD mediated inhibition of CamKIIα-positive forebrain excitatory neurons during the early postnatal window does not influence anxiety-like behavior in adulthood in CamKIIα-tTA::TRE-hM4Di bigenic male mice. Shown is a schematic (A) for the experimental paradigm for vehicle (5% sucrose) or CNO (5mg/kg) administration in the early postnatal window (P2-P14) to CamKIIα-tTA::TRE-hM4Di bigenic pups which were then assessed for anxiety-like behaviors three months post cessation of CNO treatment in adulthood in the male cohort (A). No significant difference between vehicle and PNCNO-treated male mice was noted for the percent distance travelled in the center of the open field arena (B) (n = 10 for vehicle and n = 12 for PNCNO male mice), percent distance travelled in the open arms (C) or in the closed arms (D) of the EPM (n = 14 for vehicle and n = 12 for PNCNO male mice). Results are expressed as mean ± S.E.M., and groups are compared using the two-tailed, unpaired Student’s *t*-test.

**Extended data Figure 4-1:**
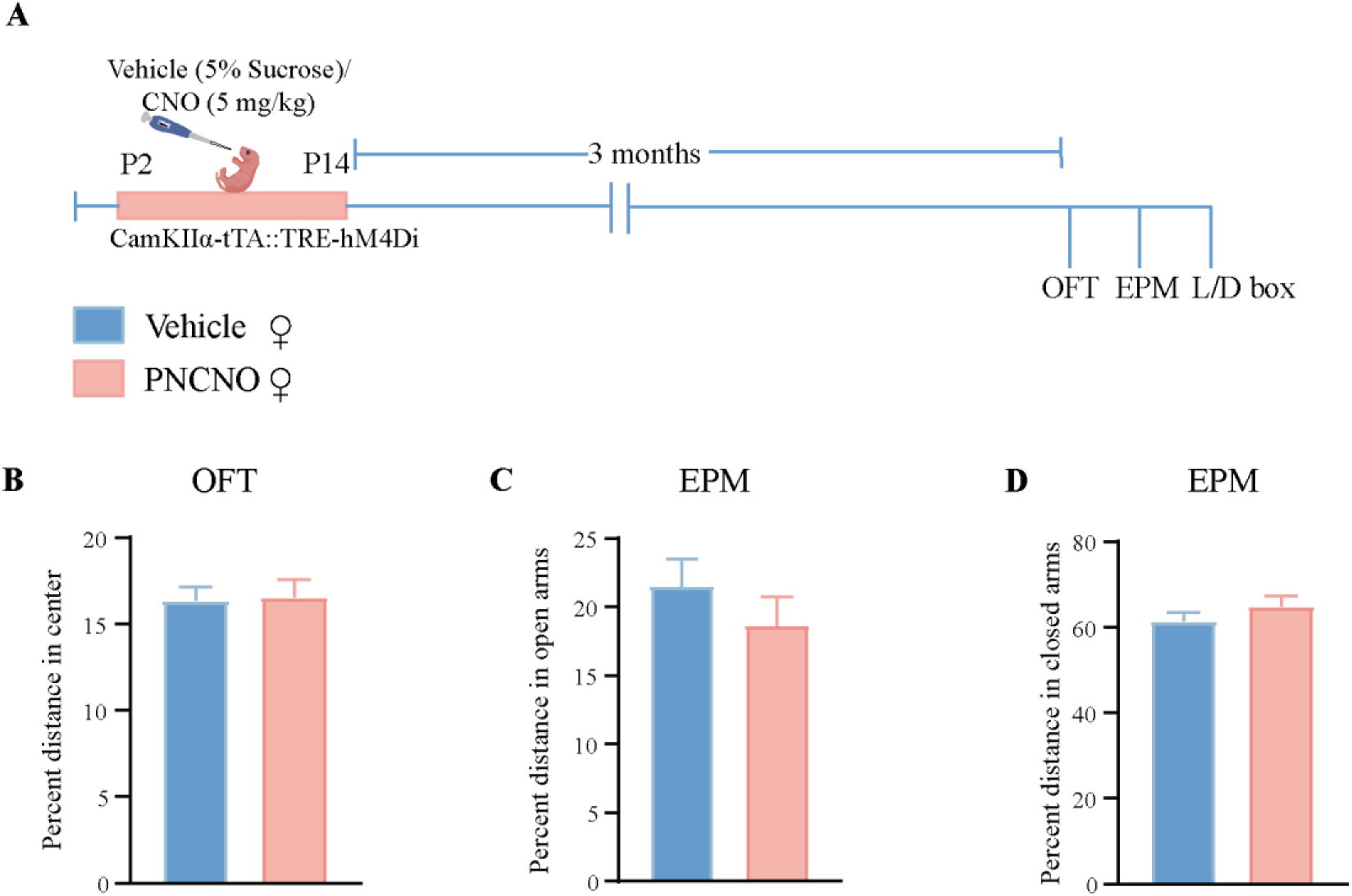
Chronic hM4Di-DREADD mediated inhibition of CamKIIα-positive forebrain excitatory neurons during the early postnatal window does not influence anxiety-like behavior in adulthood in CamKIIα-tTA::TRE-hM4Di bigenic female mice. Shown is a schematic (A) for the experimental paradigm for vehicle (5% sucrose) or CNO (5mg/kg) administration in the early postnatal window (P2-P14) to CamKIIα-tTA::TRE-hM4Di bigenic pups which were then assessed for anxiety-like behaviors three months post cessation of CNO treatment in adulthood in the female cohort (A). No significant difference between vehicle and PNCNO-treated female mice were noted for the percent distance travelled in the center of the open field box (B) (n = 10 for vehicle and n = 12 for PNCNO female mice), percent distance travelled in the open arms (C) or in the closed arms (D) of the EPM (n = 12 for vehicle and n = 11 for PNCNO female mice). Results are expressed as mean ± S.E.M., and groups are compared using the two-tailed, unpaired Student’s *t*-test.

**Extended data Figure 5-1:**
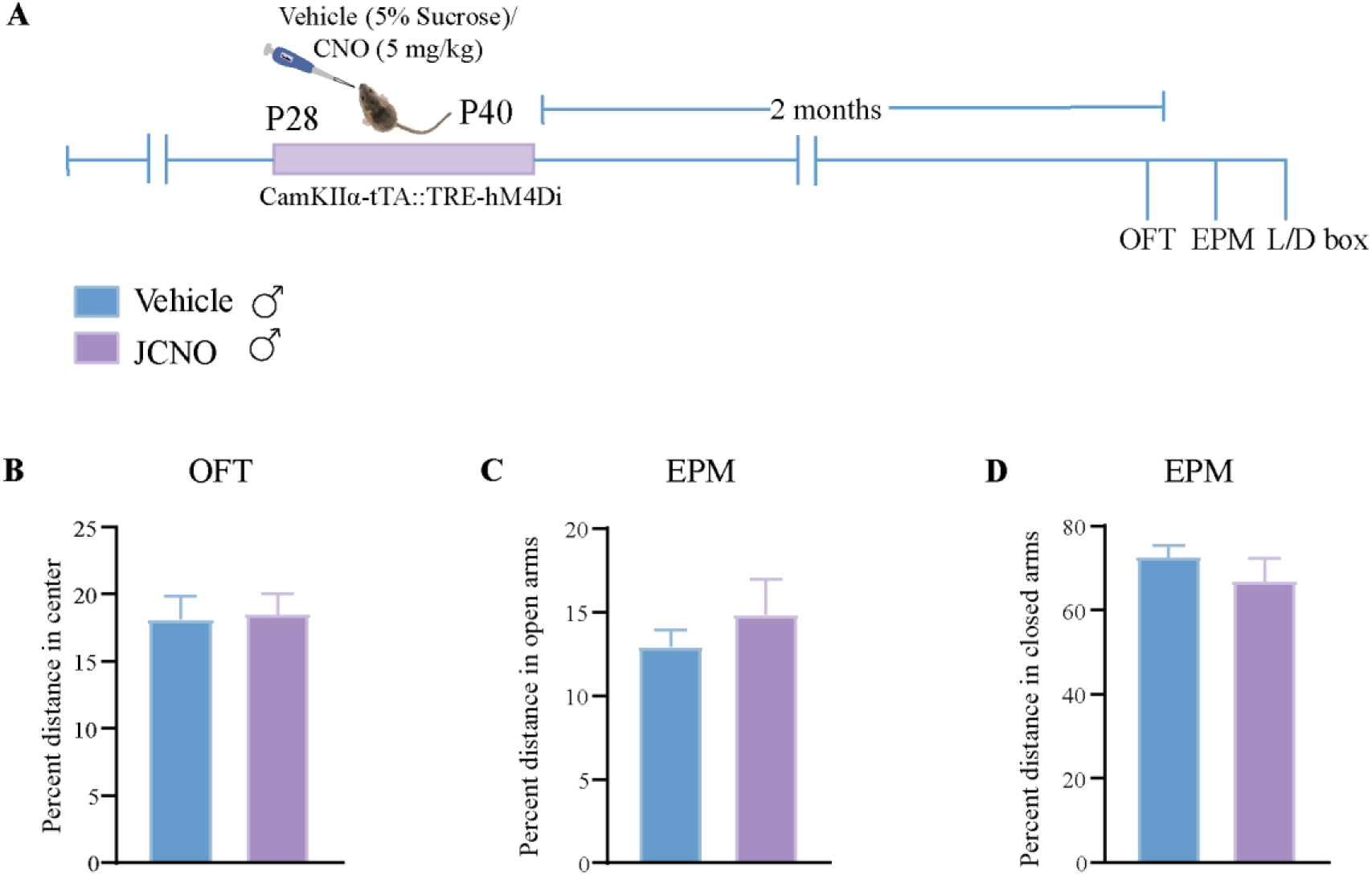
Chronic hM4Di-DREADD mediated inhibition of CamKIIα-positive forebrain excitatory neurons in the juvenile window (P28-P40) does not influence anxiety-like behavior in adulthood in CamKIIα-tTA::TRE-hM4Di bigenic male mice. Shown is a schematic (A) for the experimental paradigm for vehicle (5% sucrose) or CNO (5mg/kg) administration in the juvenile window (P28-P40) in CamKIIα-tTA::TRE-hM4Di bigenic pups which were then assessed for anxiety-like behaviors two months post cessation of CNO treatment in adulthood in the male cohort (A). No significant difference between vehicle and JCNO-treated male mice were noted for the percent distance travelled in the center of the open field box (B) (n = 18 for vehicle and n = 16 for JCNO male mice), percent distance travelled in the open arms (C) or in the closed arms (D) of the EPM (n = 18 for vehicle and n = 16 for JCNO male mice). Results are expressed as mean ± S.E.M., and groups are compared using the two-tailed, unpaired Student’s *t*-test.

## Statistics table

**Statistics table:**
| **Figure number** | **Type of test** | ***p*-value** | **t statistics** | **n (Vehicle)** | **n (CNO)** |
| --- | --- | --- | --- | --- | --- |
| **1 K** | Unpaired t-test | **0.034** | 2.905 | 4 | 3 |
| **1 L** | Unpaired t-test | **0.027** | 3.084 | 3 | 4 |
| **1 Q** | Unpaired t-test with Welch correction | 0.513 | 0.775 | 3 | 5 |
| **1 R** | Unpaired t-test | **0.018** | 3.458 | 4 | 3 |
| **2 B** | Unpaired t-test/<br>Unpaired t-test with Welch correction | P2=0.502 ,<br>P3=0.663,<br>P4=0.242,<br>P5=0.635,<br>P6=0.5449,<br>P7=0.472,<br>P8=0.473,<br>P9=0.428,<br>P10=0.583,<br>P11=0.861,<br>P12=0.738,<br>P13=0.841,<br>P14=0.973 | P2=0.699 ,<br>P3=0.449,<br>P4=1.244,<br>P5=0.501,<br>P6=0.655,<br>P7=0.770,<br>P8=0.766,<br>P9=0.826,<br>P10=0.567,<br>P11=0.179,<br>P12=0.344,<br>P13=0.206,<br>P14=0.0353 | P2=5 ,<br>P3=6,<br>P4=6,<br>P5=6,<br>P6=5,<br>P7=6,<br>P8=6,<br>P9=6,<br>P10=6,<br>P11=6,<br>P12=6,<br>P13=6,<br>P14=6 | P2=6 ,<br>P3=6,<br>P4=6,<br>P5=6,<br>P6=6,<br>P7=6,<br>P8=6,<br>P9=6,<br>P10=6,<br>P11=6,<br>P12=6,<br>P13=6,<br>P14=6 |
| **2 C** | Unpaired t-test/<br>Unpaired t-test with Welch correction | P9=0.539 , P12=0.877 | P9=0.655 ,<br>P12=0.160 | P9=6,<br>P12=5 | P9=6,<br>P12=5 |
| **2 D** | Unpaired t-test/<br>Unpaired t-test with Welch correction | P9=0.276 , P12=0.615 | P9=1.159,<br>P12=0.521 | P9=6,<br>P12=5 | P9=6,<br>P12=5 |
| **2 E** | Unpaired t-test/<br>Unpaired t-test with Welch correction | P9=0.315 , P12=0.528 | P9=1.115 ,<br>P12=0.657 | P9=6,<br>P12=5 | P9=6,<br>P12=5 |
| **2 G** | Unpaired t-test | P28=0.909 ,<br>P29=0.884,<br>P30=0.907,<br>P31=0.744,<br>P32=0.823,<br>P33=0.873,<br>P34=0.917,<br>P35=0.98,<br>P36=0.742,<br>P37=0.619,<br>P38=0.866,<br>P39=0.769,<br>P40=0.8783 | P28=0.118,<br>P29=0.15,<br>P30=0.12,<br>P31=0.337,<br>P32=0.232,<br>P33=0.164,<br>P34=0.107,<br>P35=0.026,<br>P36=0.34,<br>P37=0.518,<br>P38=0.174,<br>P39=0.304,<br>P40=0.158 | P28=5 ,<br>P29=6,<br>P30=6,<br>P31=5,<br>P32=4,<br>P33=5,<br>P34=5,<br>P35=5,<br>P36=5,<br>P37=4,<br>P38=5,<br>P39=5,<br>P40=5 | P28=5 ,<br>,<br>P29=5,<br>P30=5,<br>P31=6,<br>P32=5,<br>P33=6,<br>P34=6,<br>P35=5,<br>P36=6,<br>P37=6,<br>P38=6,<br>P39=6,<br>P40=6 |
| **3 C** | Unpaired t-test | 0.194 | 1.343 | 10 | 12 |
| **3 D** | Unpaired t-test | 0.799 | 0.258 | 10 | 12 |
| **3 E** | Unpaired t-test | 0.462 | 0.749 | 10 | 12 |
| **3 G** | Unpaired t-test | 0.399 | 0.858 | 14 | 12 |
| **3 H** | Unpaired t-test | 0.697 | 0.394 | 14 | 12 |
| **3 I** | Unpaired t-test | 0.661 | 0.444 | 14 | 12 |
| **3 J** | Unpaired t-test | 0.439 | 0.787 | 14 | 12 |
| **3 L** | Unpaired t-test with Welch correction | 0.232 | 1.246 | 14 | 12 |
| **3 M** | Unpaired t-test | 0.059 | 1.982 | 14 | 12 |
| **4 C** | Unpaired t-test | 0.127 | 1.591 | 10 | 12 |
| **4 D** | Unpaired t-test | 0.763 | 0.306 | 10 | 12 |
| **4 E** | Unpaired t-test | 0.268 | 1.14 | 10 | 12 |
| **4 G** | Unpaired t-test | 0.203 | 1.314 | 12 | 11 |
| **4 H** | Unpaired t-test | 0.402 | 0.856 | 12 | 11 |
| **4 I** | Unpaired t-test with Welch correction | 0.054 | 2.075 | 12 | 11 |
| **4 J** | Unpaired t-test | 0.232 | 1.231 | 12 | 11 |
| **4 L** | Unpaired t-test | 0.171 | 1.425 | 10 | 10 |
| **4 M** | Unpaired t-test | 0.209 | 1.302 | 10 | 10 |
| **5 C** | Unpaired t-test | 0.176 | 1.385 | 18 | 16 |
| **5 D** | Unpaired t-test | 0.7861 | 0.274 | 18 | 16 |
| **5 E** | Unpaired t-test | 0.355 | 0.939 | 18 | 16 |
| **5 G** | Unpaired t-test | 0.518 | 0.653 | 18 | 16 |
| **5 H** | Unpaired t-test with Welch correction | 0.624 | 0.497 | 18 | 16 |
| **5 I** | Unpaired t-test with Welch correction | 0.102 | 1.701 | 18 | 16 |
| **5 J** | Unpaired t-test | 0.282 | 1.094 | 18 | 16 |
| **5 L** | Unpaired t-test with Welch correction | 0.691 | 0.401 | 18 | 16 |
| **5 M** | Unpaired t-test | 0.26 | 1.147 | 18 | 16 |
| **6 B** | Unpaired t-test | 0.227 | 1.239 | 14 | 12 |
| **6 C** | Unpaired t-test | 0.539 | 0.624 | 14 | 12 |
| **6 E** | Unpaired t-test | 0.802 | 0.255 | 10 | 10 |
| **6 F** | Unpaired t-test | 0.471 | 0.736 | 10 | 10 |
| **6 H** | Unpaired t-test | 0.495 | 0.695 | 11 | 11 |
| **6 I** | Unpaired t-test | 0.553 | 0.603 | 11 | 11 |
| **7 C** | Unpaired t-test | **0.018** | 2.596 | 10 | 10 |
| **7 D** | Unpaired t-test | 69dB(+4)=0.401,<br>73dB(+8)=0.310,<br>81dB(+16)=0.138 | 69dB(+4)=0.86,<br>73dB(+8)=1.044,<br>81dB(+16)=1.554 | 10,<br>10,<br>10 | 10,<br>10,<br>10 |
| **7 F** | Unpaired t-test | 0.968 | 0.04 | 11 | 9 |
| **7 G** | Unpaired<br>t-test | 69dB(+4)=0.333,<br>73dB(+8)=0.798,<br>81dB(+16)=0.698 | 69dB(+4)=0.994,<br>73dB(+8)=0.26,<br>81dB(+16)=0.394 | 11,<br>11,<br>11 | 9,<br>9,<br>9 |
| **Extended**<br>**Data Fig**<br>**3-1 B** | Unpaired<br>t-test | 0.747 | 0.328 | 10 | 12 |
| **Extended**<br>**Data Fig**<br>**3-1 C** | Unpaired<br>t-test | 0.988 | 0.015 | 14 | 12 |
| **Extended**<br>**Data Fig**<br>**3-1 D** | Unpaired<br>t-test with<br>Welch<br>correction | 0.756 | 0.315 | 14 | 12 |
| **Extended**<br>**Data Fig**<br>**4-1 B** | Unpaired<br>t-test | 0.882 | 0.151 | 10 | 12 |
| **Extended**<br>**Data Fig**<br>**4-1 C** | Unpaired<br>t-test | 0.328 | 1.003 | 12 | 11 |
| **Extended**<br>**Data Fig**<br>**4-1 D** | Unpaired<br>t-test | 0.307 | 1.046 | 12 | 11 |
| **Extended**<br>**Data Fig**<br>**5-1 B** | Unpaired<br>t-test | 0.864 | 0.173 | 18 | 16 |
| **Extended**<br>**Data Fig**<br>**5-1 C** | Unpaired<br>t-test with<br>Welch<br>correction | 0.425 | 0.814 | 18 | 16 |
| **Extended**<br>**Data Fig**<br>**5-1 D** | Unpaired<br>t-test with<br>Welch<br>correction | 0.354 | 0.947 | 18 | 16 |

